# The unique contributions of Rab11 and Rab35 to the completion of cell division

**DOI:** 10.1101/2024.09.25.614863

**Authors:** Paulius Gibieža, Emilija Ratkevičiūtė, Girstautė Dabkevičiūtė, Vilma Petrikaitė

## Abstract

Rab11 and Rab35 were shown to be responsible for wide-scale intracellular membrane trafficking during the last phases of the cell cycle. Even though data suggests that Rab11 and Rab35 are abscission-related, each gives a different phenotype to dividing cells. Moreover, nobody has ever looked at both and systematically compared the two during the regulation of cancer cell division. Hence, the molecular interplay and compensatory mechanisms between Rab11 and Rab35 remain to be established. Our data shows that depletion of Rab11 or Rab35 inhibits cytokinetic abscission and is associated with aberrant levels of F-actin at the intercellular bridge. In contrast, overexpression of functionally similar Rab protein cannot rescue downregulation-associated cytokinetic defects in cancer cells. We also demonstrate that even though Rab11 and Rab35 function in the same molecular pathway, the input that each individual Rab11 and Rab35 carries out during a distinct mitotic M-phase stage differs. Thus, this research study is dedicated to getting a more in-depth understanding of molecular machinery behind Rab11- and Rab35-mediated cell progression from late anaphase to cytokinesis.

## Introduction

Cytokinesis occurs at the end of mitosis, when the cytoplasm that connects two newly formed daughter cells is cleaved in two, separating the cells (Fededa and Gerlich, 2012; Schiel and Prekeris, 2013; Carlton et al., 2020). For the successful division to occur, cells must experience a tremendous number of cytoskeletal modifications and highly coordinated intracellular processes, which would allow for smooth and equivalent distribution of cellular compartments between the two, resulting in a physical separation of dividing cells. The actual cessation of the newly forming cells starts in late anaphase when cell cytoplasm undergoes contraction via the formation of an actomyosin ring made of filamentous actin (F-actin), motor protein myosin-II, formins and many actin cross-linkers (Pollard, 2010; Chew et al., 2017; Pollard and O’Shaughnessy, 2019). The actomyosin-based contractile ring forms under the plasma membrane’s surface and is positioned so that it can separate chromosomes. However, the constriction of the plasma membrane alone is insufficient to separate newly formed cells. When the contractile ring constricts to its absolute capacity, it forms an intercellular bridge (ICB), which now possesses large amounts of F-actin and a mass of highly compacted central spindle microtubule bundles.

For quite a while, several researchers have been trying to clarify the regulation of molecular machinery that drives the removal of actin deposits and central spindle microtubules from the ICB during cytokinesis. Here, Rab GTPases were shown to deliver specialized membranous vesicles and various proteins mediating cytokinesis (Yu et al., 2007; Miserey-Lenkei and Colombo, 2016; Nakayama, 2016). The considerable attention that this topic has drawn emphasizes the importance of this process to cell life and homeostasis as a unit of the whole system, which, if misregulated, can lead to various diseases, including cancer (Saito et al., 2004; Nguyen et al., 2009; Dong and Wu, 2018). In particular, this is relevant to the Rab GTPases-mediated cytokinesis event, which, upon dysregulation, can cause aneuploidy and multinucleation, which eventually induce tumorigenesis (Fujiwara et al., 2005; Högnäs et al., 2012; Guadagno and Progida, 2019). In principle, successful cytokinesis requires spatial and temporal membrane trafficking, which Rab11 and Rab35 primarily regulate from the entire Rab protein family. These molecular switches belong to a family of small monomeric GTPases. When GTP-bound via its effectors FIP3 and FIP4, Rab11 targets intracellular vesicles derived from recycling endosomes along the central spindle microtubules to the cleavage furrow during late cytokinesis (Fielding et al., 2005, p. 3; Wilson et al., 2005). Normally, Rab11-containing endosomes are distributed throughout the cytoplasm during metaphase and anaphase. As the mitotic phase progresses towards the end, these Rab11 endosomes are concentrated at the cleavage furrow and regulate telophase and final cytokinetic division in mammalian cells (Wilson et al., 2005). This mediates cortical actin depolymerization at the abscission site by recruiting and transporting SCAMP2/3 and p50RhoGAP to the ICB. This inhibits RhoGTPase activity and reduces F-actin polymerization, narrowing the ICB (Schiel et al., 2012; Gibieža and Petrikaitė, 2021). Additionally, Rab11-FIP3 and FIP4 complexes were shown to interact with Arf6 and regulate membrane trafficking to the sites of the ICB and the midbody (MB) before the final abscission (Fielding et al., 2005, p. 3; Takahashi et al., 2011).

Like Rab11, Rab35 is also responsible for F-actin clearance from the abscission site. Rab35-containing endosomes are found throughout the cytoplasm during interphase and accumulate at the ingressing furrow and ICB during early and late cytokinesis, respectively (Kouranti et al., 2006). It lowers the amount of phosphatidylinositol-4,5-biphosphate (PtdIns(4,5)P_2_), which otherwise would maintain the stability of the existing ICB via the promotion of actin polymerization (Dambournet et al., 2011). Rab35 recruits its effector, Oculo-Cerebro-Renal syndrome of Lowe (OCRL) lipid phosphatase, to stop this process. It targets OCRL to the ICB, where local hydrolysis of PtdIns(4,5)P_2_ finally occurs. In this way, and because of the reduced cortical F-actin oligomerization, the F-actin levels at the ICB are kept at a level sufficient for normal cytokinetic abscission to occur (Kouranti et al., 2006; Dambournet et al., 2011, p. 35). Additionally, Rab35 recruits another effector, MICAL1 oxidoreductase, which catalyzes the oxidation of the two methionine residues of F-actin (M44 and M47). It thus activates the depolymerization of actin filaments at the abscission site (Hung et al., 2011; Frémont et al., 2017b; Niu et al., 2020), which narrows the ICB to such an extent that it is required for the incoming microtubule-severing enzyme spastin to bind to CHMP1B. Then, this molecular interaction activates the ESCRT-III machinery, targeted to the MB, to coordinate the mitotic spindle disassembly (Connell et al., 2009; Caballe and Martin-Serrano, 2011; Vietri et al., 2015). CHMP4B assembles with the ESCRT-III subunit at the end of cytokinetic abscission and forms the cortical membrane-distorting helical filaments. This process mediates the complete termination of the remaining ICB at the secondary ingression site. It physically separates newly formed daughter cells (Carlton and Martin-Serrano, 2007; Morita et al., 2007; Elia et al., 2011; Christ et al., 2016).

Plenty of data is available, indicating that depending on a research object, type of species, tissue type, cell line, and many cellular factors, either overexpression or downregulation of Rab11 or Rab35 can induce cancer. Indeed, one of the studies showed that Rab11a is highly expressed in hepatocellular carcinoma (HCC) and is associated with poor clinical prognosis (Zhang et al., 2020). In this particular study, overexpression of Rab11a promoted proliferation, migration, and invasion of human HCC cell lines *in vitro*. In contrast, the downregulation of Rab11a inhibited those tumorigenesis-associated processes. Similarly to these findings, Rab11a positively influenced the growth of HCC cells in vivo (Zhang et al.. Another study by *Wang* et al. reported that Rab11-family interacting proteins (Rab11-FIPs) is overexpressed in human colorectal cancer (CRC), which directly correlated to poor patient prognosis (Wang et al., 2018). Indeed, increased expression of Rab11-FIP4 was linked to induced proliferative capacity, migration, invasion and tumorigenic potential of tumour cells *in vivo* (Wang et al., 2018). At the same time, a study by *Ye* et al. demonstrated that as part of the Grb2-DENND1A-Rab35 signalling cascade, Rab35 could regulate gastric cancer cell migration and invasion (Ye et al., 2018). Moreover, the phenotype of reduced cell adhesion and induced cell migration was determined after the downregulation of Rab35. Indeed, these cellular properties were closely related to epithelial-mesenchymal transition (Allaire et al., 2013). Moreover, it was shown that the dysregulation of Rab11 and Rab35 in different disease contexts could be an oncogenic factor (Gopal Krishnan et al., 2020).

Despite all the data available, the literature lacks information about the specific phenotypic differences that Rab11 and Rab35 grant to dividing cancer cells. Also, nobody looked at the possible interplay between these two proteins while regulating processes necessary to cells, especially during cell division. The only attempt to demonstrate the interaction between Rab11 and Rab35 was by *Iannantuono* et al., where Rab11-FIP1 was shown to maintain Rab35 at the ICB to mediate actin removal during abscission. In contrast, the absence of Rab11-FIP1 was demonstrated to reduce Rab35 and further cytokinetic defects (Iannantuono and Emery, 2021). Additionally, since these proteins play functionally similar roles during cell division (Chesneau et al., 2012; Gibieža and Petrikaitė, 2021), it would be interesting to see whether the downregulation of each or both at the same time, in terms of the regulation of cytokinesis, would have any effect on the progression of cancer. Another intriguing point would be determining whether downregulation-associated phenotypic defects in cancer cells could be rescued by overexpressing the same or functionally similar gene.

For the reasons stated above, we used an RNAi approach to downregulate an individual and different combinations of Rab11a, Rab11b, and Rab35 genes and performed subsequent rescue experiments to characterize the phenotype of dividing cervical cancer cells.

## Materials and Methods

### Cell culture and transfections

The human adenocarcinoma HeLa-wt cell line was obtained from the American Type Culture Collection (ATCC, Manassas, USA). HeLa-wt cells were maintained in 5% CO_2_ and at 37°C in Dulbecco’s modified Eagle’s medium (DMEM, Life Technologies Gibco, USA), supplemented with 10% fetal bovine serum (Life Technologies Gibco, USA) and a 1% solution of 10,000 units/ml penicillin and 10 mg/ml streptomycin (SIGMA, USA). Cells were regularly tested for mycoplasma contamination. For rescuing with Rab11a and Rab35 downregulation-associated phenotypic alterations in HeLa-wt cells, siRNA-resistant Rab11a-GFP or Rab35-GFP plasmids were transfected into siRNA-treated cells using Lipofectamine 2000 (Invitrogen, USA, #11668019), according to the manufacturer’s instruction.

### Plasmids

pEGFP-C1-Rab11a wt and pEGFP-C1-Rab35 wt plasmids were received from Prof. Rytis Prekeris, University of Colorado Anschutz Medical Campus, Denver, USA.

### Antibodies and other staining reagents

The following primary antibodies were used for immunofluorescence and Western blots: acetylated-α-tubulin (Cell Signalling, USA, #D20G3, IF-1:200), anti-Rab11a (Sigma Life Science, USA, #HPA051697, WB-1:250), anti-Rab11b (Sigma Life Science, USA, #HPA054396, WB-1:250), anti-Rab35 (Abcam, USA, #ab300116, WB-1:250), beta-tubulin loading control (Invitrogen Thermo Fisher Scientific, USA, #MA5-16308, WB-1:2000). The following secondary antibodies were used for immunofluorescence: Alexa Flour 488 donkey anti-rabbit IgG (H+L) (Thermo Fisher Scientific, USA, #A21206, IF-1:250).

The staining dyes used for immunofluorescence include Alexa Flour phalloidin 568 (Thermo Fisher Scientific, USA, #A12380, IF-1:50 from a 40× stock) and DAPI (Thermo Fisher Scientific, USA, #D1306, IF-1 µg/ml).

### Immunofluorescence

For immunofluorescence analysis, cells plated on collagen I-coated (at final 50 µg/ml concentration, Life Technologies Gibco, USA) No. 1 microscope cover glasses (VWR) were fixed using 4% paraformaldehyde (PFA) for 15 min at room temperature (R.T.). The cover glasses were then washed thrice with 1× PBS and permeabilized with 1× PBS / 0.2% Triton X-100 (Thermo Fisher Scientific, USA, #A160046.AE) permeabilization buffer for 3 min at R.T. Following permeabilization, the cells were washed three times with 1× PBS / 1% bovine serum albumin (BSA) wash buffer. Next, the cover glasses were incubated with primary antibodies premixed in a wash buffer for 1 hour at 37 °C and a moisture-maintaining incubator, followed by three 5-min-long antibody washing steps with a wash buffer. Next, the cover glasses were incubated with secondary antibodies and prepared in a wash buffer for 30 minutes at 37 °C, in a moisture-maintained incubator. Then, the antibody was rinsed off while washing the cells thrice for 5 min each, using a wash buffer. Finally, the cover glasses were incubated with 1 µg/ml DAPI (Thermo Fisher Scientific, USA, #D1306) solution for 10 min at R.T., followed by a final 3-round washing step with a wash buffer before being mounted on microscopic slides with a Prolong Diamond Antifade mounting media (Invitrogen, USA, #P36965). The next day, immunofluorescence was detected using an inverted fluorescence Olympus IX73 microscope (Olympus Europe Holding Gmbh) using a 20× lens for multinucleation and mitotic stage analysis assays, and upright confocal Olympus Fluoview FV1000 microscope (Olympus Europe Holding Gmbh) using either a 20×, 40× and 60× oil lens for actin fluorescence intensity measurements, depending on the requirements. In total, three biological repeats were completed, each containing from 5 to 10 technical replicates.

### Western blotting

For Western blotting, cells were scraped in 1× PBS containing 1 mM PMSF (Thermo Fisher Scientific, USA, #36978) and 1% Triton X-100 (Thermo Fisher Scientific, USA, #A160046.AE) and incubated for 30 min on ice. The collected cells were centrifuged at 15,000 g for 5 min at 4 °C to pellet the lysates. The total protein was determined using a Bradford assay reagent (Thermo Fisher Scientific, Pierce, USA, #23238). Samples were mixed with 5× SDS sample loading buffer (Thermo Fisher Scientific, Pierce, USA, #39000) and boiled for 5 min at 95 °C. Depending on their molecular weight, the boiled samples were loaded onto SDS-PAGE gel (12%) to separate proteins. Proteins were then transferred onto PVDF membranes and stained with the indicated antibodies under standard laboratory procedures. Beta tubulin loading control was used, and the relative expression of the proteins in each sample was normalized to the level of tubulin. The amount of protein was quantified by densitometry using Image J software (https://imagej.nih.gov/ij/). A total of 3 biological repeats were performed, and the data presented is an average.

### Multinucleation assay

HeLa-wt cells transfected with non-targeting siRNA (control) were seeded on collagen I-coated (at final 50 μg/ml concentration, Life Technologies Gibco, USA) No. 1 microscope cover glasses (VWR, USA), which after 72 h in culture were fixed in 4% PFA (Thermo Fisher Scientific, USA). For various siRNA-treated cells, 24 h post-seeding on collagen I-coated glass coverslips, HeLa-wt cells were treated with appropriate siRNA constructs. This allowed for 48 h for siRNA expression, followed by a fixation as previously described. Fixed cells were permeabilized with 0.2% Triton X-100 (Thermo Fisher Scientific, USA, #A160046.AE) for 3 min at R.T. and stained with Alexa Flour 568 phalloidin (Thermo Fisher Scientific, USA, #A12380) for 30 min at 37°C and with 1 µg/ml DAPI (Thermo Fisher Scientific, USA, #D1306) for 10 min at R.T. The cover glasses were mounted on glass microscope slides using Prolong Diamond Antifade mountant (Invitrogen, USA, #P36965). Then, random fields on the coverslips were photographed using an inverted fluorescence Olympus IX73 microscope (Olympus Europe Holding Gmbh) with a 20× lens. The number of multinucleated cells was counted manually, including binucleated and poly-lobed cells and micronuclei. The rate of total multinucleated cells was then calculated in ten randomly chosen fields for each biological repeat. All cells were valued as one population. All data are derived from at least three independent experiments (biological repeats), each containing 5-15 technical replicates.

To rescue multinucleation, different siRNA constructs together with respective GFP-tagged Rab11a-or Rab35-coding plasmids were transfected into HeLa-wt cells seeded on collagen I-coated No. 1 microscope cover glasses (VWR, USA) according to Lipofectamine 2000 manufacturer’s (Invitrogen, USA, #11668019) instruction. Then, 48 h post-transfection, cells were processed and examined for multinucleation as described above.

### Mitotic stage analysis

To determine the percentage fraction of cells in the telophase phase, HeLa-wt cells transfected with non-targeting siRNA (control) or with various siRNA-treated cells and with either Rab11a-GFP or Rab35-GFP rescued cells were fixed and stained with a combination of anti-acetylated-α-tubulin antibody (Cell Signalling, USA, #D20G3, IF-1:200) and DAPI (Thermo Fisher Scientific, USA, #D1306), as described above. Cells in telophase were identified based on a mitotic spindle and chromatin condensation status. Cells were counted in 10 randomly selected fields in each independent experiment and expressed as a percentage of all cells. All data are derived from at least three independent experiments (biological replicates), where each replicate contains 5-15 technical replicates.

### Actin fluorescence intensity measurements

The cells in late telophase, which had entirely reformed their cytoskeleton and no longer possessed cleavage furrow, were selected for this experiment. A single-image plane with a cell in late telophase, which includes the entire ICB in focus, was selected to measure F-actin fluorescence at the ICB. The regions of interest (ROIs) were then selected at the ICB to analyze F-actin enrichment. For the control (non-targeting siRNA), ROIs were selected at the opposing poles of the cell. Fluorescence intensity was then measured as a sum-fluorescence and normalized to the ROI size. Lastly, the ratio of fluorescence in the ICB and opposing poles was calculated to measure the enrichment of the F-actin at the ICB. The enrichment of actin fluorescence was counted in 10 carefully selected cytokinetic ICBs in each independent experiment and expressed as a percentage change compared to the control. All data are derived from at least three independent experiments (biological repeats), with ten technical replicates for each biological repeat.

### siRNA treatments

Using different siRNAs, Lipofectamine 2000 (Invitrogen, USA, #11668019) transfected HeLa-wt cells. Double-stranded interfering RNA sequences used for siRNA treatment are as given: Silencer Select Negative Control #1 siRNA (Ambion, USA, #4390843); Rab11a - 5’-CAACAAUGUGGUUCCUAUUtt-3’ (Ambion, USA #4390824); Rab11b - 5’-CUAACGUAGAGGAAGCAUUtt-3’ (Ambion, USA #4390824); Rab35 - 5’-GCAGUUUACUGUUGCGUUUtt-3’ (Ambion, USA #4390824). The transfection of single siRNA or a combination of different siRNAs and the rescue of siRNA-treated cells with co-transfection of plasmid DNA and siRNA were completed according to the manufacturer’s (Invitrogen) recommendations. Each siRNA transfection experiment was performed at least three times (biological repeats), with three technical repeats.

### Statistical analysis

All statistical analyses were performed using Microsoft Office Excel data analysis tool. A Student’s *t*-test was performed on all datasets to determine the significance level if otherwise noted. Whenever three or more groups were compared, One-way ANOVA was used to perform statistical analysis. For multinucleation assay, mitotic stage analysis, and actin fluorescence intensity measurement, at least five randomly chosen image fields from a single coverslip were used for data collection. The experiments were repeated three or more times (technical replicates). All data are derived from at least three independent experiments (biological replicates). The same exposure was used for all the specimens in that experiment to quantify immunofluorescence, and the data from images was quantified using ImageJ software (National Institutes of Health, USA). All bars in the graphs show mean ± S.D. unless otherwise indicated.

## Results

### The interconnected expression of intracellular Rab11 and Rab35 influences cytokinesis

First, we used the RNA interference (RNAi) approach to downregulate Rab11a, Rab11b and Rab35 using different siRNAs. As expected, having almost 90% sequence identity, the downregulation of Rab11b led to a reduction in Rab11a protein level by 28,1%, which is shown by W.B. analysis (Fig 1A). On the other hand, upon downregulation of Rab11a, the intracellular levels of Rab11b decreased by 42,1% compared to the total protein levels of Rab11b in the control (Fig 1B). This data thus indicates that the protein levels of different Rab11 isoforms are interrelated *in vitro*, where the reduced expression of Rab11a affects the expression of Rab11b and vice versa. It is worth mentioning that the relative level of Rab11a did not change and was equal to 100,9% after the downregulation of Rab35 (Fig 1A). Contrary to Rab11a, the protein level of Rab11b decreased to 29,8% after the downregulation of Rab35 (Fig 1B). Meanwhile, upon the downregulation of Rab11a, the protein level of Rab35 was about 18,5% (Fig 1C). Surprisingly, it increased to 108,2% after the treatment with Rab11b siRNA compared to the control (Fig 1C).

**Figure 1.**
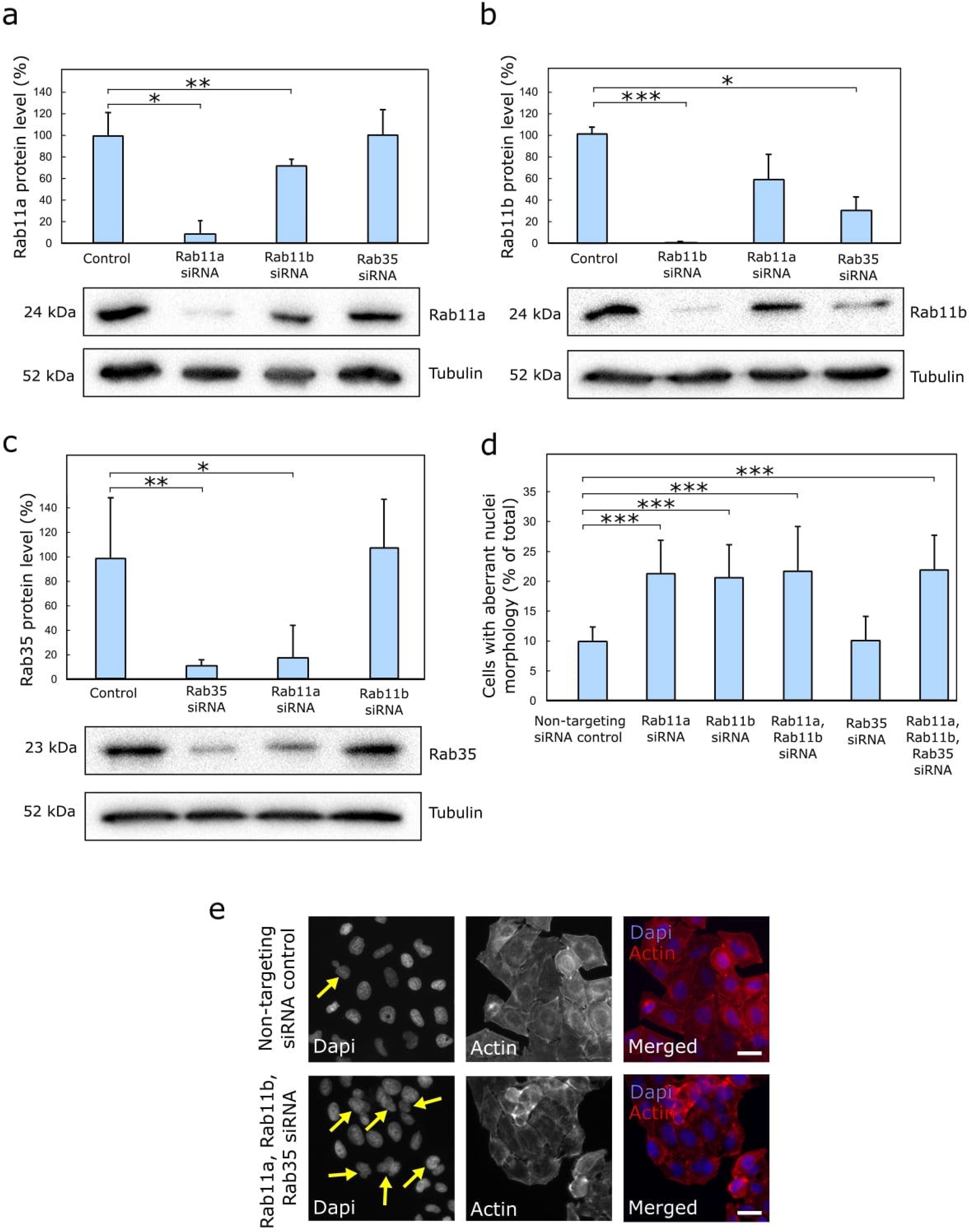
Downregulation of Rab11a, Rab11b and Rab35 induces multinucleation, including binucleation, poly-lobed nuclei and micronuclei. **(A-C)** HeLa-wt cells transfected with different siRNA were cultured until confluency. Lysates were then collected, and the relative protein levels of either Rab11a, Rab11b or Rab35 were measured using Western blot analysis with the indicated antibodies. Quantifications of the relative protein levels in Rab11a-siRNA-, Rab11b-siRNA- and Rab35-siRNA-treated cells are presented above the W.B. blots. Beta tubulin was used as a loading control, and the relative expression of the proteins was normalized to the level of intracellular tubulin. The data shown are the means and S.D. derived from at least three independent experiments. * - labels statistically significant differences (p < 0.05). **(D)** Control and different siRNA-treated HeLa-wt cells were fixed stained with DAPI and phalloidin, and the combined number of binucleated, poly-lobed and micronuclei-containing cells was assessed. * - labels statistically significant differences (p < 0.05). The data shown are the means and S.D. derived from at least three independent experiments. At least 3000 individual cells were captured using 20× magnification for each study group unless otherwise stated **(E)** The images represent the population of HeLa-wt cells treated with non-targeting siRNA control and with combined siRNA-treated cells that contain binucleated, poly-lobed and micronuclei containing cells (scale bar represents 20 µm). Arrows indicate binucleated, poly-lobed, and micronuclei-posing cells formed by cytokinesis failure. Unless otherwise stated, at least 30 individual images were captured using 20× magnification for each study group. n=3.

The data above suggests that Rab11a, Rab11b and Rab35 function in the same molecular pathway, where Rab35 functions downstream of Rab11a, as the downregulation of Rab35 does not affect Rab11a level (Fig 1A). In contrast, downregulation of Rab11a reduces the level of Rab35 (Fig 1C). On the contrary, Rab35 functions upstream of Rab11b because the downregulation of Rab35 reduces the level of Rab11b (Fig 1B). In contrast, the downregulation of Rab11b does not reduce the level of Rab35 (Fig 1C). The level of Rab35 upon downregulation of Rab11b even increases by 8,2% when compared to the control (Fig 1C), which indicates that cells attempt to compensate for the lost function and, therefore, increase the production of Rab35, which could then possibly take over the function of Rab11b. If needed, the expression levels and localization patterns of Rab11 and Rab35 under normal conditions and upon downregulation can be found in the following research articles (Fielding et al., 2005; Wilson et al., 2005; Kouranti et al., 2006; Klinkert et al., 2016).

Next, we fixed with non-targeting siRNA control and different siRNA-transfected HeLa-wt cells and stained them with phalloidin and DAPI. This allowed us to visualize cell nuclei in blue and separate cells from one another by labelling cells’ cortex with red actin stain - phalloidin 568 (Fig 1E). Under normal physiological conditions, the HeLa-wt population contains ∼5% abnormal mitotic and multinucleate cells *in vitro* (Yasuda et al., 2006). To compare, in our study, 9,9% of the entire population of HeLa-wt cells treated with non-targeting siRNA control contained abnormal nuclei morphology (Fig 1D). Next, we found that downregulation of either Rab11a or Rab11b leads to significantly abnormal nuclei morphology, where 21,3% and 20,6% of the entire population contained multinucleated cells, respectively (Fig 1D). Meanwhile, knocking down Rab11a and Rab11b simultaneously increased this abnormal morphology to 21,8% (Fig 1D). In addition, after a combined K.D. with Rab11a, Rab11b and Rab35 siRNAs, this number increased to 22,0% (Fig 1D-E). This was over two times higher than the control. It is worth noting that siRNA-resistant transfection with a plasmid coding for GFP-Rab11a fully rescued with Rab11a downregulation-associated cytokinetic defects in cancer cells (Supplemental Fig S1A). Surprisingly, cytokinetic defects of Rab11a-siRNA treated cells could also be rescued by overexpressing siRNA-resistant GFP-Rab35 (Supplemental Fig S1A). This data goes in hand with W.B. results and strengthens the proposal that Rab11a and Rab35 function in the same molecular pathway, where Rab35 functions downstream of Rab11a, as the rescue effect on Rab11a-downregulated cells by GFP-Rab11a is more substantial than compared to the rescue effect by GFP-Rab35 (Supplemental Fig S1A). This is the first indication of the interplay between Rab11 and Rab35 during the regulation of cancer cell division. Meanwhile, as the downregulation of Rab35 via RNAi did not affect nuclei morphology (Fig 1D), the rescue results with siRNA-resistant GFP-Rab35 and GFP-Rab11a plasmids become negligible (Supplemental Fig S1B).

Additionally, we performed a W.B. analysis on individual Rab proteins to exclude the possibility of the side effects of double or even triple K.D. treatment, which could potentially affect the expression of a protein of interest. We measured the expression profile of each Rab protein under a combined double or triple downregulation. According to the results, none of the experimental conditions impacted the expression level of each Rab protein. The K.D. still proved efficient in causing phenotypic alterations in cancer cells (Supplemental Fig S3A-C).

### Rab11 and Rab35 remove actin from the intercellular bridge and complete cytokinesis

Next, we fixed cells to check the number of cells stuck in the telophase phase. This was followed by staining with phalloidin 568 (F-actin marker) and anti-acetylated-α-tubulin antibody (marks ICB). This combination of dyes allowed us to visualize cells in the telophase phase with ICB formed in the middle of the dividing cells (Fig 2B and Supplemental Fig S2C). We found that amongst all siRNA-treated cells, the downregulation of Rab35 had the highest impact on the number of cells in the telophase phase. Here, in comparison to 0,8% of the control (Fig 2A-B), 3,5% of the entire Rab35 siRNA-treated population were in the telophase phase (Fig 2A) at the scope, which is also apparent from the fluorescent images (Supplemental Fig S2C). Unfortunately, the difference between the number of cells in the telophase phase between the control and the remaining study groups was less significant (Fig 2A).

**Figure 2.**
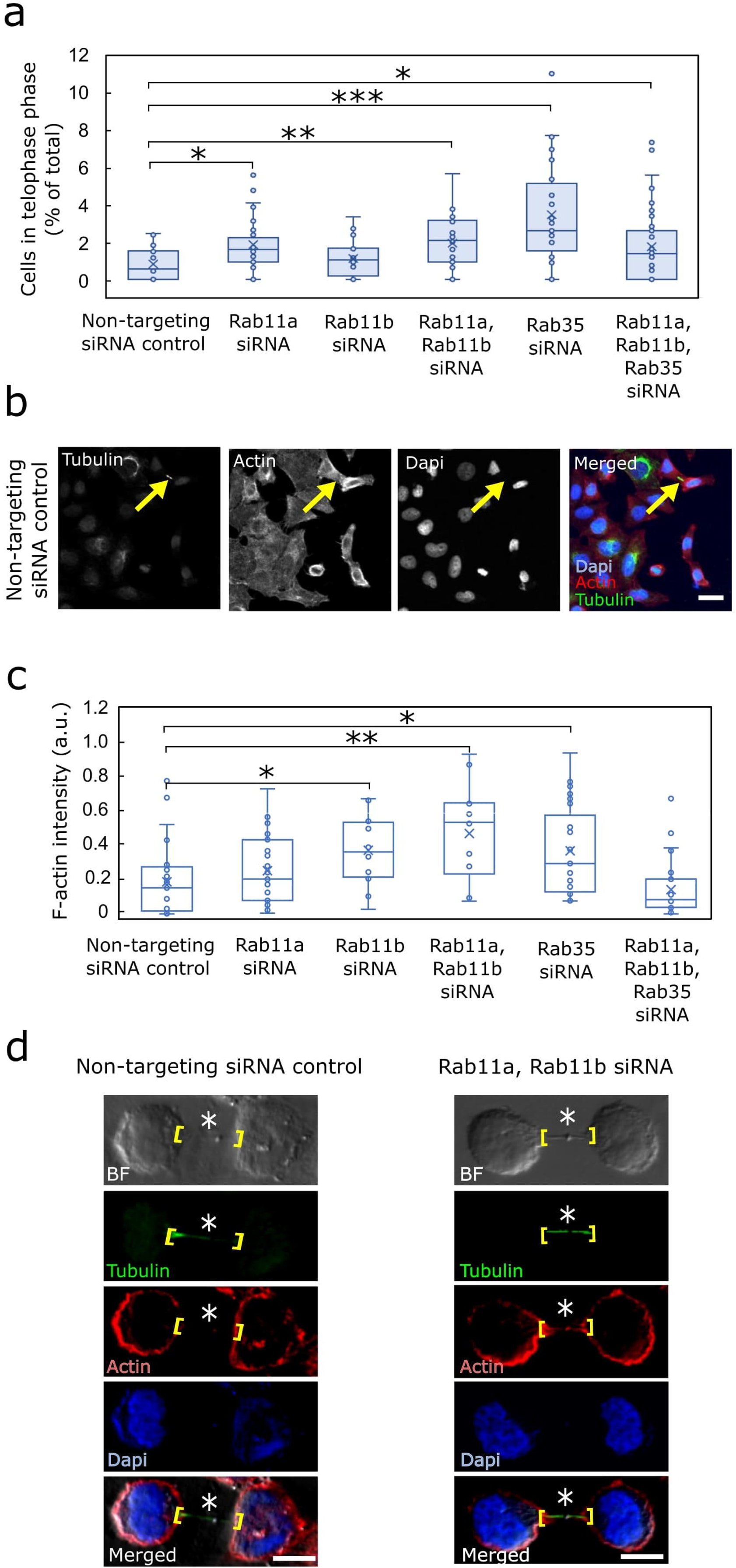
Downregulation of different Rabs increases the number of cells in the telophase phase, leading to elevated levels of F-actin at the intercellular bridge. **(A)** HeLa-wt cells were transfected with different siRNAs, fixed, and stained with DAPI, phalloidin and anti-acetylated-α-tubulin primary antibody. Then, the number of cells in telophase was counted manually. * - labels statistically significant differences (p < 0.05). Unless otherwise stated, at least 30 different images (at least 3000 cells) with varying numbers of cells were inspected using 20× magnification for each study group. n=3. **(B)** The images represent non-targeting siRNA control-treated HeLa-wt cells in telophase, identified by the ICB labelled with anti-acetylated-α-tubulin primary antibody (scale bar indicates 20 µm). Arrows in the images point to the ICB, which had formed during cytokinesis. Unless otherwise stated, at least 30 individual images (at least 3000 cells) were captured using 20× magnification for each study group. n=3. **(C)** Control cells were processed as specified in Fig 2A. Then, actin enrichment at the ICB was evaluated by confocal microscopy using a 60× magnification oil lens. The quantification of actin level at the ICB in control vs. individual siRNA-treated cells, or a combination of different siRNAs, was performed. * - labels statistically significant differences (p < 0.05). At least 30 different cells were evaluated for each study group. n=3 **(D)** siRNA-treated cells that could not remove the accumulated actin possessed an increased level of actin at their ICB compared to the control (scale bar indicates 10 µm). The site where the ICB was formed and where the F-actin levels were measured are shown in brackets. Asterisks mark the midbody. Unless otherwise stated, at least 30 individual cells using 60× magnification were assessed for each study group. n=3.

Another important aspect of this assay to be tested was rescuing the phenotypic defects by overexpressing different Rabs. According to the results, neither the rescue with the same gene that had been downregulated nor the overexpression of a functionally similar gene showed a statistically significant rescue effect to the number of cells in the telophase phase for both Rab11a-or Rab35-siRNA treated cells (Supplemental Fig S2A-B). Remarkably, only the rescue of Rab35-depleted cells with siRNA-resistant GFP-Rab35 was the closest to reducing the number of cells in the telophase phase resembling the control (Supplemental Fig S2B).

Hereafter, we measured the filamentous actin (F-actin) level at the ICB during late telophase, an important factor limiting the cells’ ability to separate, as accumulated actin impedes smooth ICB abscission. The late ICBs can be easily spotted; when the ICB is at its last stage, just before the final abscission step, it becomes long and narrow (Fig 2D, brightfield and tubulin-labelled images). It can be easily cut. To measure the F-actin level and compare between different study groups, the control and various siRNA-treated cells were prepared for examination with confocal microscopy, as described in a previous section. A high magnification (60× oil lens) confocal microscopy was used to image the late ICB between the dividing cells. To evaluate the F-actin level at the ICB, a corrected total cell fluorescence (CTCF) (Jakic et al., 2017) was counted, values normalized to the background, and the ratio between the F-actin enrichment at the ICB versus the cell pole was evaluated.

According to the results, the accumulated F-actin levels were determined separately in Rab11a and Rab11b siRNA-treated cells and upon simultaneous downregulation of both Rab11 isoforms, - Rab11a and Rab11b (Fig 2C). Here, the normalized F-actin levels at the furrow reached 0,26 (relative units), 0,37 (relative units) and 0,47 (relative units), respectively, and were higher than 0,24 (relative units) in the control cells treated with non-targeting siRNA (Fig 2C). A similar accumulation of F-actin at the ICB between the newly forming daughter cells was also identified in Rab35 siRNA-treated cells (Fig 2C). The enrichment of F-actin in this study group reached 0,37 (relative units) and was 54,2% higher than in the control.

The results above indicate that Rab11 and Rab35 are essential regulators of cytoskeleton modifications during the last phases of cell division. Significantly, the dysregulation of this process may disturb actin removal from the ICB and potentially disrupt the cleavage of the ICB at the abscission site during cytokinesis. Because of this mitotic exit problem, cells experience morphological alterations in their nuclei, leading to polyploidy and tetraploidy, which, if uncontrolled, can later develop into tumorigenic phenotypes (Fig 3) (Fujiwara et al., 2005).

**Figure 3.**
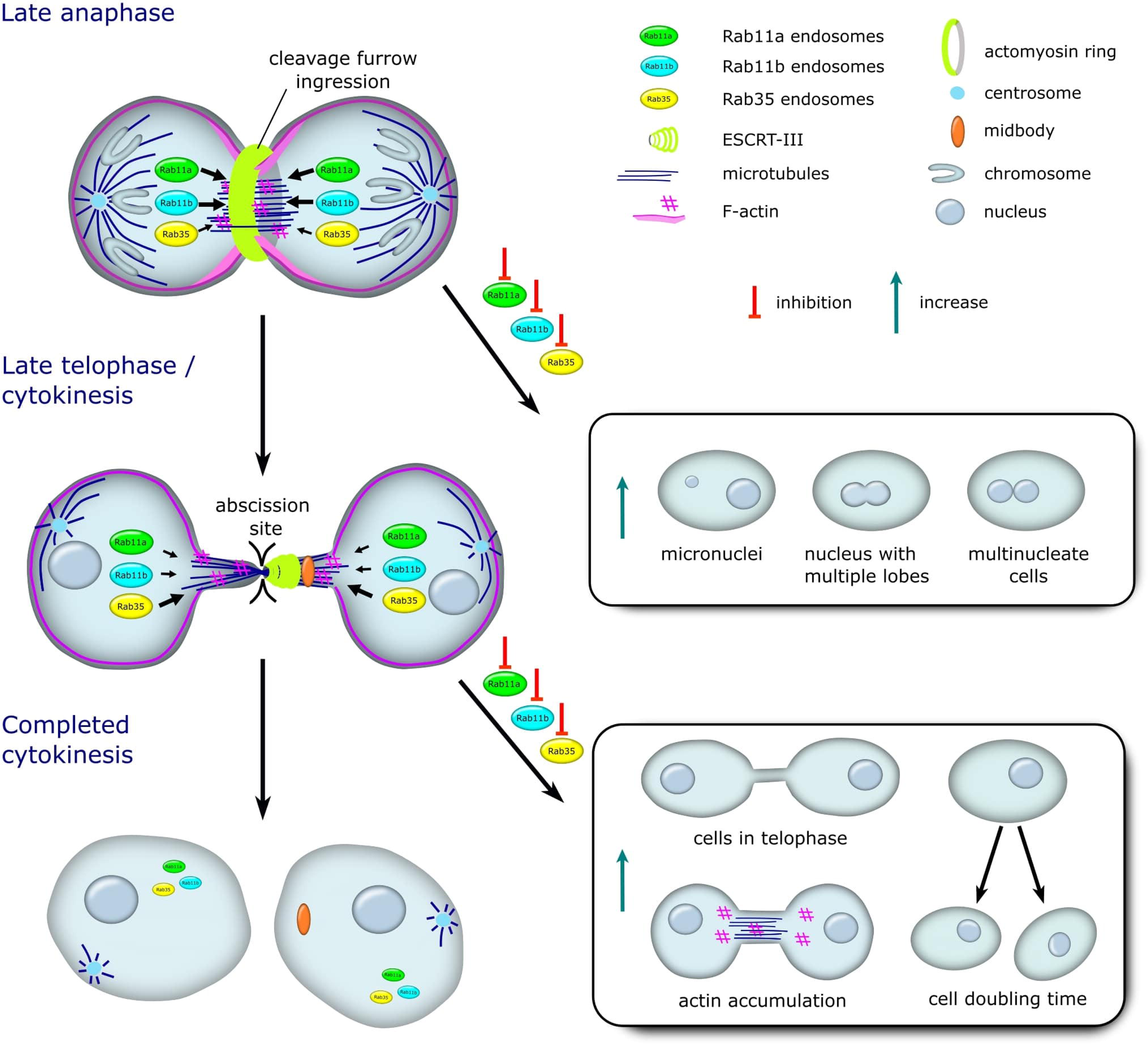
A model representing an input that each Rab11 and Rab35 plays throughout the transition from late anaphase to the completed cytokinesis. On the left side of the image, the size of the arrows next to the endosome’s depiction correlates to the relevance and input of the specific Rab protein to a particular phase of the cell cycle. The right side of the image shows the processes that appear because of Rab protein inhibition during the formation of the cleavage furrow ingression and the generation of the abscission site.

## Discussion

Cell division is a complex and highly regulated event where cellular cytoskeleton remodelling and intracellular transport events must work in concert for the newly formed daughter cells to accomplish cell division and to separate into two distinct cells physically. Specifically, Rab11 and Rab35 were shown to mediate the endocytic transport to the division site in cells which are preparing to complete division or are at the actual physical division process (Frémont et al., 2017a; Laflamme et al., 2017).

In this study, we found out that the downregulation of individual isoforms of Rab11 influences the protein level of other Rab11 isoforms still expressed by the cells (Fig 1A-B). Since Rab11 isoforms share high sequence homology, these results were as expected (Vale-Costa and Amorim, 2016). Additionally, upon the depletion of Rab11b, cells try to compensate for the lost function and increase the production of Rab35 instead (Fig 1C). As Rab11 and Rab35 are both abscission-related, this gave us a first glimpse of the potential interplay between the two. Moreover, our results suggested that both isoforms of Rab11 and Rab35 function in the same molecular pathway. Indeed, Rab35 functions downstream of Rab11a but upstream of Rab11b (Fig 1A-C).

The recycling endosomes Rab11 and Rab35 emerged as the key regulators of specialized disassembly of locally accumulated F-actin network at the ICB during cytokinesis (Gibieža and Petrikaitė, 2021). So, we decided to test how the downregulation of an individual or a combination of different Rabs may affect the cleavage furrow formation during late anaphase (Fig 3). The downregulation of either Rab11a or Rab11b led to a significant increase in multinucleated cells, where more than 20% of cells of the entire population had an aberrant nuclei morphology (Fig 1D). To compare, *Yu* et al. took a different approach, and by transfecting HeLa cells with a dominant-negative mutant of Rab11, they demonstrated that multinucleation increases to around 15% (Yu et al., 2007). In addition, the depletion of individual Rab11a or Rab11b by siRNA led to around 13% formation in binuclear cells (Yu et al., 2007). In another study, a defective function of Rab11 in *Drosophila* spermatocytes led to a faulty actin constriction ring and ingression furrow formation (Giansanti et al., 2008). However, in our study, adding up the effects of Rab11a and Rab11b co-depletion did not significantly increase with downregulation-associated phenotypic defects during cleavage furrow formation (Fig 1D).

Meanwhile, upon the downregulation of Rab35, the formation of multinucleated cells did not increase. It stayed at the control level of around 10% (Fig 1D). In a similar approach using *Drosophila* as a study model, a downregulation of Rab35 via RNAi determined that binucleation increases to around 5% and 20%, 3 and 6 days post-transfection, respectively (Kouranti et al., 2006). For comparison, our results were obtained three days after the transfection with the corresponding siRNA. Unfortunately, after making a simultaneous triple-KD with Rab11a, Rab11b and Rab35 siRNAs, the phenotypic defects to the cleavage furrow formation did not add up, where only 22,0% of cells had an aberrant nuclei morphology (Fig 1D). Surprisingly, downregulation-associated phenotypic defects during the cleavage furrow formation could be restored by overexpressing the same gene or the gene of another functionally similar Rab protein (Supplemental Fig S1A-B). Once again, this proposes the presence of molecular interplay between Rab11 and Rab35.

Next, we decided to study whether the downregulation of Rab proteins affects cells’ ability to proceed from late telophase to cytokinesis, which predetermines whether the cell would divide successfully or not. To this end, the number of cells in the telophase phase was evaluated after downregulating Rab proteins. According to the results, only the role of Rab35 became protuberant, as the depletion of Rab35 increased the number of cells in telophase to 3,5% - more than four times as much as in the non-targeting siRNA control-treated cells (Fig 2A). This means Rab35 is less relevant to the cleavage furrow formation but is more important for setting up an abscission site at the ICB (Fig 3). Compared with Dambournet et al., it was demonstrated that the K.D. of Rab35 increases the time required for the final abscission, which directly predetermined the fatal fate of the dividing cells (Dambournet et al., 2011). Finally, neither an individual nor a combined downregulation of the rest of the Rabs in this study showed a significant difference, as the telophase cells were just around the 2% level mark (Fig 2A).

There is a requirement for disassembling this rigid ICB, as cells cannot get released from one another. Here, many different proteins come into action. However, the ESCRT-III complex remains one of the leading players, activating the spastin-binding cascade and dismantling microtubule bundles that constitute the ICB (Goliand et al., 2018). Meanwhile, the F-actin at the ICB is best disassembled by the complex action from Rab35 and Rab11 endosomes. Indeed, Rab35 binds to MICAL1 and OCRL and recruits them to the F-actin depolymerization process. At the same time, Rab11 delivers p50RhoGAP and thus reduces F-actin polymerization at the ICB (Gibieža and Petrikaitė, 2021). Activating the later effectors helps to reduce F-actin levels at the ICB to a point where the ESCRT-III could come in and make a final cut (Goliand et al., 2018) (Fig 3).

Thus, consistent with mitotic stage analysis data, it is evident that Rab35 is essential for setting up the abscission site and removing the accumulated F-actin from the ICB (Fig 2A and Supplemental Fig S2B-C). To highlight the significance of Rab35 in the completion of cytokinesis, a study by *Dambournet* et al. showed that cells depleted either for Rab35 or its effector OCRL are characterized by an increased accumulation of PtdIns(4,5)P_2_ and F-actin at the ICB (Dambournet et al., 2011). Consequently, it causes abscission defects in cells, which can be restored by treating the cells with F-actin depolymerizing drugs, such as Latrunculin A (Dambournet et al., 2011). In most studies that try to elucidate the mechanisms governing cellular division, the inability of cells to depolymerize F-actin from the abscission site at the ICB seems to be the main reason for unsuccessful cellular division. For example, in one of our studies, we demonstrated that another Rab’s family member, Rab14, is also involved in actin removal from the abscission site. A Rab14-knockout disrupted cytokinesis and increased F-actin levels at the mitotic ICB (Gibieža et al., 2021). Consistent with the literature, in this study, we showed that cells individually depleted for Rab11a, Rab11b and Rab35 or co-depleted for Rab11a and Rab11b struggled to remove the F-actin from the mitotic ICB (Fig 2C-D).

### Conclusions

To summarize all the results, Rab11 isoforms are interrelated in cells because the reduced expression of Rab11a influences the expression level of Rab11b and vice versa. We also showed that upon the depletion of one Rab protein, cells try to compensate for the lost function and thus increase the production of another Rab protein of a similar function. Moreover, we demonstrated that Rab11 and Rab35 function in the same molecular pathway, where Rab35 function downstream of Rab11a but upstream of Rab11b. Even though both Rab11 and Rab35 were shown to be essential regulators of cell division, we clarified the different inputs each protein has during different phases of the mitotic division. Both Rab11 isoforms are more important for setting up the cleavage furrow ingression during late anaphase. At the same time, Rab35 may function as the primary regulator of an abscission site formation during the late telophase/cytokinesis phase. According to the results, the cytokinetic defects caused by the downregulation of Rab11 or Rab35 are associated with abnormal levels of F-actin at the ICB. Importantly, albeit intracellular protein levels vary during gene manipulation analysis, simultaneous depletion of Rab11 and Rab35 had no additive effect on cytokinesis-related defects. Hence, the mechanism for molecular interplay is not entirely proven and requires further examination.

## Data availability

The data generated in this study are available upon request from the corresponding author.

## Acknowledgements

We thank Prof. Rytis Prekeris (University of Colorado Anschutz Medical Campus) for the pEGFP-C1-Rab11a wt and pEGFP-C1-Rab35 wt plasmids.

## Authors’ Contributions

Conceptualization, P.G.; Methodology, P.G., E.R. and G.D.; Validation, P.G.; Formal analysis, P.G., E.R. and G.D.; Investigation, P.G.; Data curation, P.G.; Writing - original draft preparation, P.G.; Writing - review and editing, P.G., E.R., G.D. and V.P.; Visualization, V.P.; Supervision, P.G. and V.P.; Project administration, P.G.; Funding acquisition, P.G. and V.M. All authors have read and agreed to the published version of the manuscript.

## Supplementary Figure legends

**Supplemental Figure 1. Downregulation of Rab11 and Rab35 promotes multinucleation, including forming poly-lobed and binucleated nuclei and micronuclei.**

**(A-B)** Non-targeting siRNA control and different siRNA-treated HeLa-wt cells were fixed stained with DAPI and phalloidin, and the successful cellular division was measured by assessing the combined number of multinucleate cells, including binucleated, poly-lobed and micronuclei containing cells. The number of cells was expressed as the percentage of the total number. The image includes the rescue of K.D. with the same gene or overexpression of a different gene. * - labels statistically significant differences (p < 0.05). The data shown are the means and SD derived from at least three independent experiments. Unless otherwise stated, at least 30 images (at least 3000 cells) with varying numbers of cells were captured using 20× magnification for each study group.

**Supplemental Figure 2. Downregulation of Rab11 and Rab35 leads to increased accumulation of F-actin at the intercellular bridge.**

**(A-B)** HeLa-wt cells seeded on collagen-coated coverslips were transfected with different siRNA or a combination of siRNAs, fixed, and stained with DAPI, phalloidin and anti-acetylated-α-tubulin primary antibody. Then, the number of cells in the telophase phase was counted manually. The image includes the rescue of K.D. with the same gene or overexpression of a different gene. * - labels statistically significant differences (p < 0.05). Data shown are the means and S.D. derived from at least three independent experiments. Unless otherwise stated, at least 30 images (at least 3000 cells) with varying numbers of cells were captured using 20× magnification for each study group. **(C)** The images represent the increased number of cells in the telophase phase after the treatment with Rab35 siRNA. This was identified by labelling the ICB with an anti-acetylated-α-tubulin primary antibody. Arrows in the images point to different ICBs formed during late telophase (scale bar indicates 20 µm).

**Supplemental Figure 3. Neither double nor triple knockdown of Rab proteins affects the individual expression of Rab11a, Rab11b and Rab35.**

**(A-C)** HeLa-wt cells transfected with individual siRNA or a combination of different siRNAs were cultured until confluency. Lysates were then collected, and the relative protein levels of either Rab11a, Rab11b or Rab35 were measured using Western blot analysis with the indicated antibodies. Quantifications of the relative protein levels of Rab11a, Rab11b and Rab35 are presented above the W.B. blots. Beta tubulin was used as a loading control, and the relative expression of the proteins was normalized to the level of intracellular tubulin.

