## Supplementary figures and images for "The unique contributions of Rab11 and Rab35 to the completion of cell division"

### Supplemental Figure 1

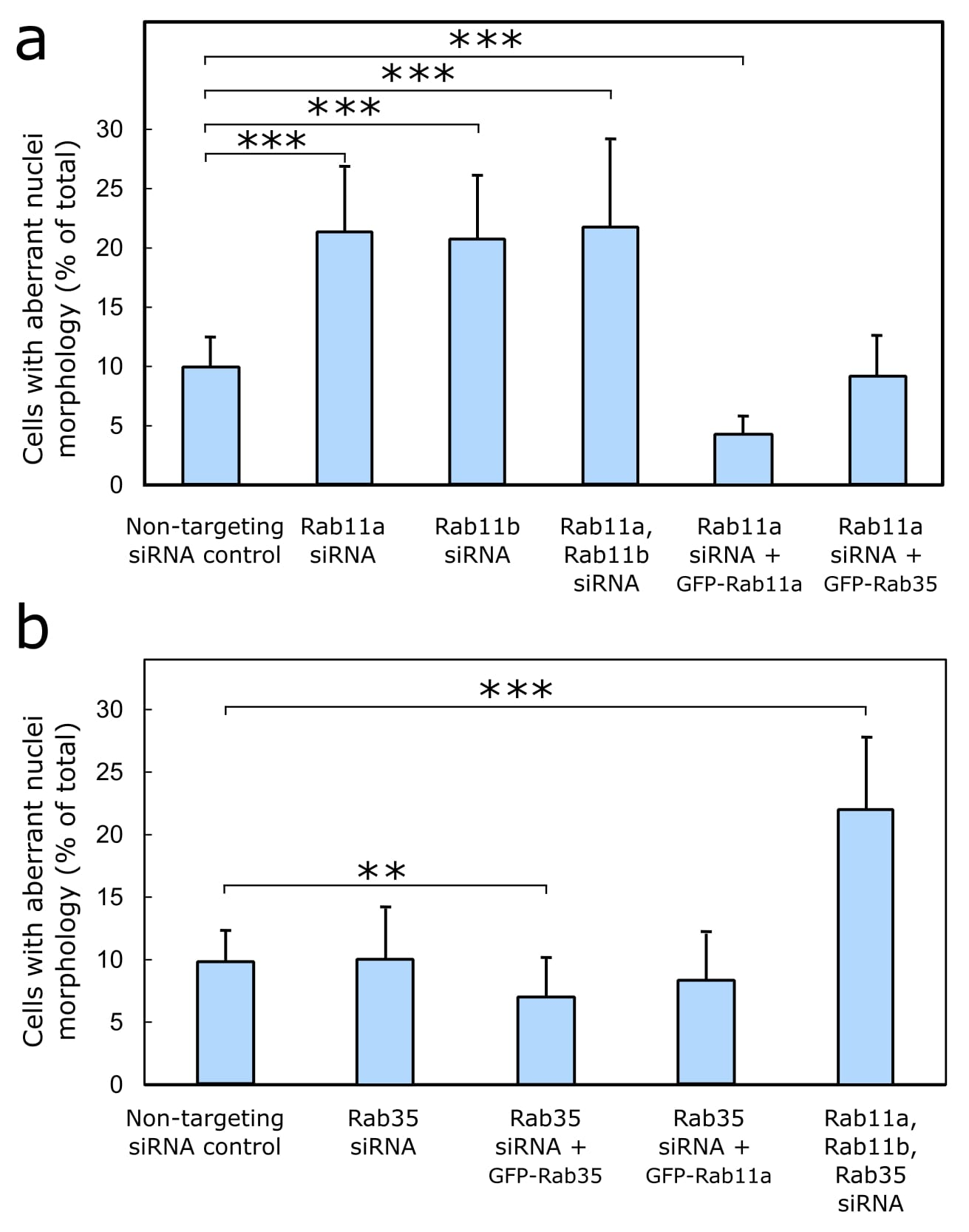

### Supplemental Figure 2

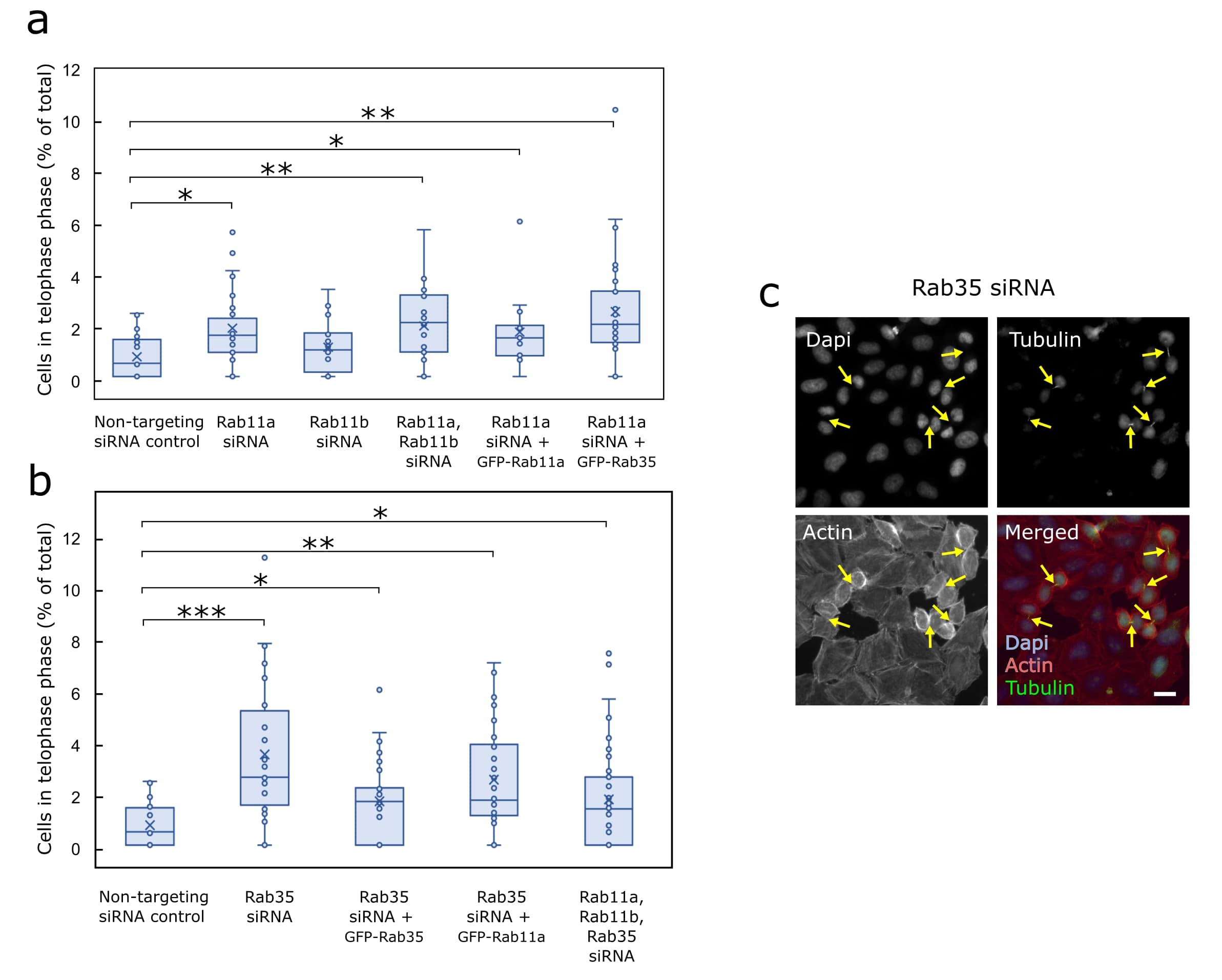

### Supplemental Figure 3

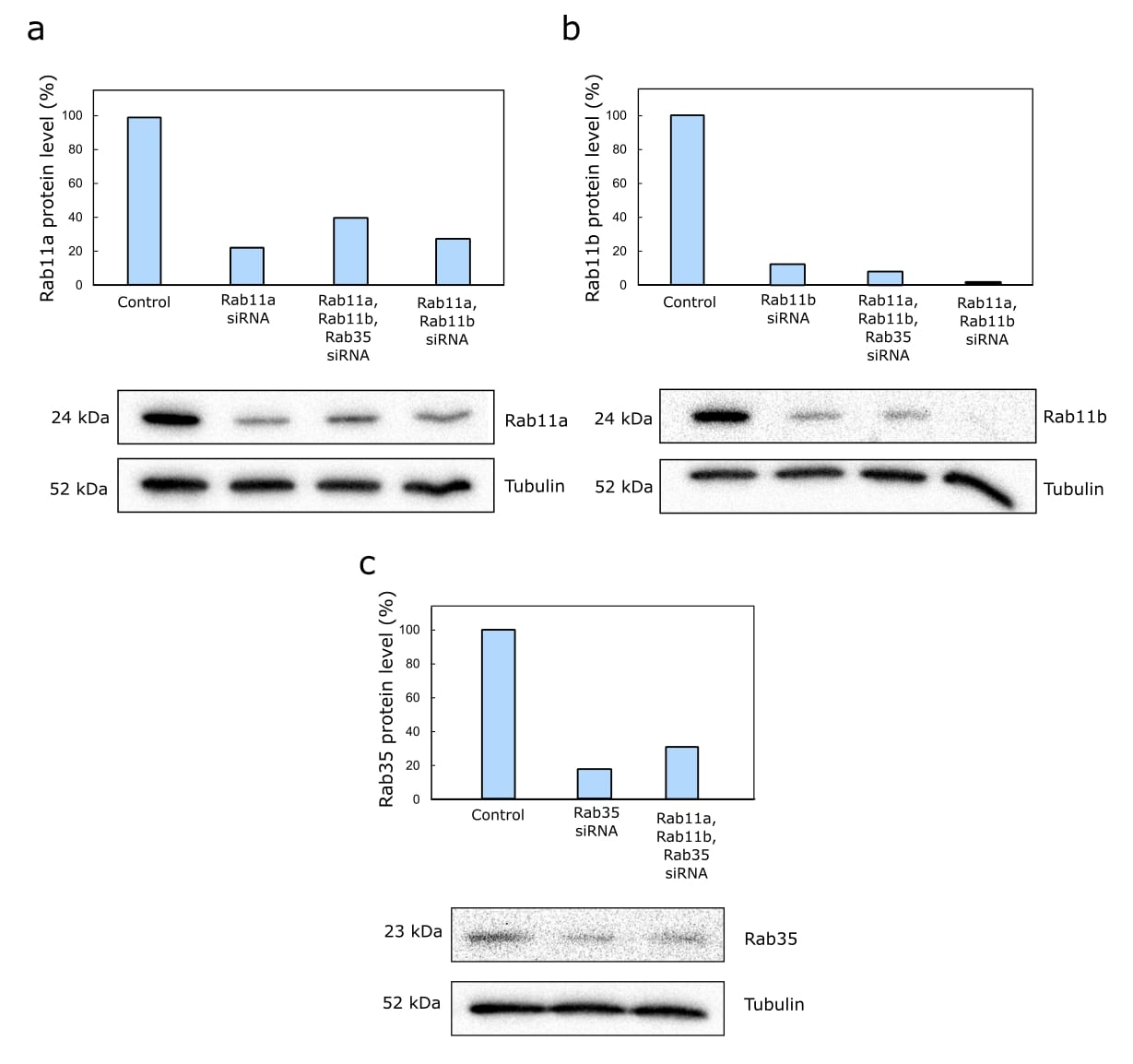
